# Optogenetic perturbations reveal temporal integration of Wnt signaling during pattern formation

**DOI:** 10.64898/2026.07.08.737164

**Authors:** Sera L. Weevers, Nicole Repina, Panagiotis Papasaikas, Jacqueline Ferralli, Sebastien Smallwood, Jörg Wittlieb, Alexander Klimovich, Charisios D. Tsiairis

## Abstract

The Wnt signaling pathway is a conserved regulator of tissue patterning and regeneration, yet how cells interpret dynamic Wnt inputs to generate robust developmental outcomes remains poorly understood. Here, we establish the first optogenetically activatable *Hydra* line, enabling precise temporal control of canonical Wnt signaling *in vivo*. Optogenetic stimulation induced dose-dependent patterning phenotypes whose rate of progression scaled with stimulation intensity. Transcriptomic analysis revealed that distinct combinations of signal intensity and duration converged onto shared transcriptional trajectories and could be described by an effective exposure metric that represents the cumulative signaling input. Functionally, Wnt activation rescued head regeneration under conditions that normally prevent organizer formation, and equivalent regenerative outcomes could be achieved through either strong, short-lived stimulation or weaker, prolonged activation. Together, our results indicate that *Hydra* tissues decode Wnt signaling through temporal integration of cumulative pathway activity, progressively accumulating transcriptional responses until patterning thresholds are reached. These findings establish a quantitative framework for understanding how dynamic morphogen signaling is translated into stable developmental decisions during regeneration.

## Introduction

Canonical Wnt signaling is a deeply conserved developmental signaling pathway that regulates cell fate specification, tissue organization, and stem cell maintenance across metazoans (1). In its canonical form, Wnt ligand binding to Frizzled and LRP co-receptors inhibits the β-catenin destruction complex, allowing β-catenin to accumulate and translocate into the nucleus, where it regulates transcription together with TCF transcription factors (2). Through this mechanism, Wnt signaling controls diverse developmental and homeostatic processes ranging from axis formation to tissue regeneration (3, 4). In many developmental contexts, Wnt proteins function as morphogens, instructing cell fate decisions through dynamically regulated signaling gradients distributed across space and time (5, 6).

During development, morphogenetic signaling pathways rarely operate as static inputs (7). Instead, signaling activity often varies dynamically in space and time, raising a fundamental question in developmental biology: how do cells interpret signaling dynamics to generate robust and reproducible patterning outcomes? Increasing evidence from multiple systems suggests that cells can respond not only to signal amplitude, but also to the duration, timing, and cumulative history of pathway activation (8). However, it remains unclear how dynamic signaling inputs are integrated to guide robust pattern formation.

The freshwater cnidarian *Hydra* provides a powerful system for studying how Wnt signaling controls axial patterning and regeneration (9). *Hydra* possesses a simple body plan organized along an oral–aboral axis, consisting of a head region surrounded by tentacles at the oral end and a basal disc at the aboral end. Canonical Wnt signaling is highest in the hypostomal organizer region (10) where graded pathway activity is associated with oral fate specification and axial organization (11), and is both necessary and sufficient for head formation (12, 13). During regeneration, injury-induced Wnt activation marks the future site of organizer formation and is required for successful axis re-establishment (14). Despite extensive work defining the role of Wnt signaling in *Hydra* patterning, it remains unclear how the dynamics of pathway activation are interpreted to drive cell fate changes.

Optogenetics provides a powerful approach to address such questions, as it enables signaling pathways to be manipulated with high spatial and temporal precision (15-17). Due to its ability to undergo reversible oligomerization upon illumination, the blue-light responsive *Arabidopsis thaliana* cryptochrome 2 (Cry2) protein is among the most widely used optogenetic tools for controlling intracellular signaling pathways (18). One such system, optoWnt, consists of Cry2 fused to the cytoplasmic domain of the Wnt co-receptor LRP6 (19). Light-induced Cry2 oligomerization promotes the clustering of LRP6 domains into signalosomes, resulting in inhibition of the destruction complex, and activation of canonical Wnt signaling (20).

Here, we establish the first optogenetically activatable *Hydra* line and use it to investigate how dynamic Wnt signaling is decoded during pattern formation and regeneration. By systematically varying the duration and intensity of optogenetic stimulation, we find that tissues respond primarily to cumulative Wnt exposure rather than instantaneous pathway activity. Distinct stimulation regimes converge onto similar transcriptional states and can generate similar regenerative outcomes through compensatory combinations of signal strength and duration. Together, our findings demonstrate that organizer formation depends on the temporal integration of Wnt signaling and establish a quantitative framework for understanding how dynamic morphogen inputs are converted into stable patterning decisions.

## Results

### Establishment of an optogenetically activatable Wnt signaling system in *Hydra*

To enable precise temporal control of canonical Wnt signaling in *Hydra*, we optimized the optoWnt system previously developed in mammalian cells for expression in *Hydra* cells (Fig. 1A) (19). To achieve this, we introduced two modifications to the original construct (Fig. 1B, Suppl. Fig 1). First, the coding sequence was codon-optimized according to the AT-rich codon usage preference of *Hydra* genes. Second, the N-terminal sequence of the *Hydra* actin gene (*HyAct*) was fused upstream of Cry2, following established transgenic design principles in *Hydra*. The resulting construct was introduced into embryos of the *Hydra vulgaris* AEP strain, and transgenic hatchlings were selected based on RFP expression (Fig. 1C). Through clonal propagation, we established stable optoWnt lines with either bilateral epithelial expression or ectoderm-restricted expression (Fig. 1D).

**Figure 1.**
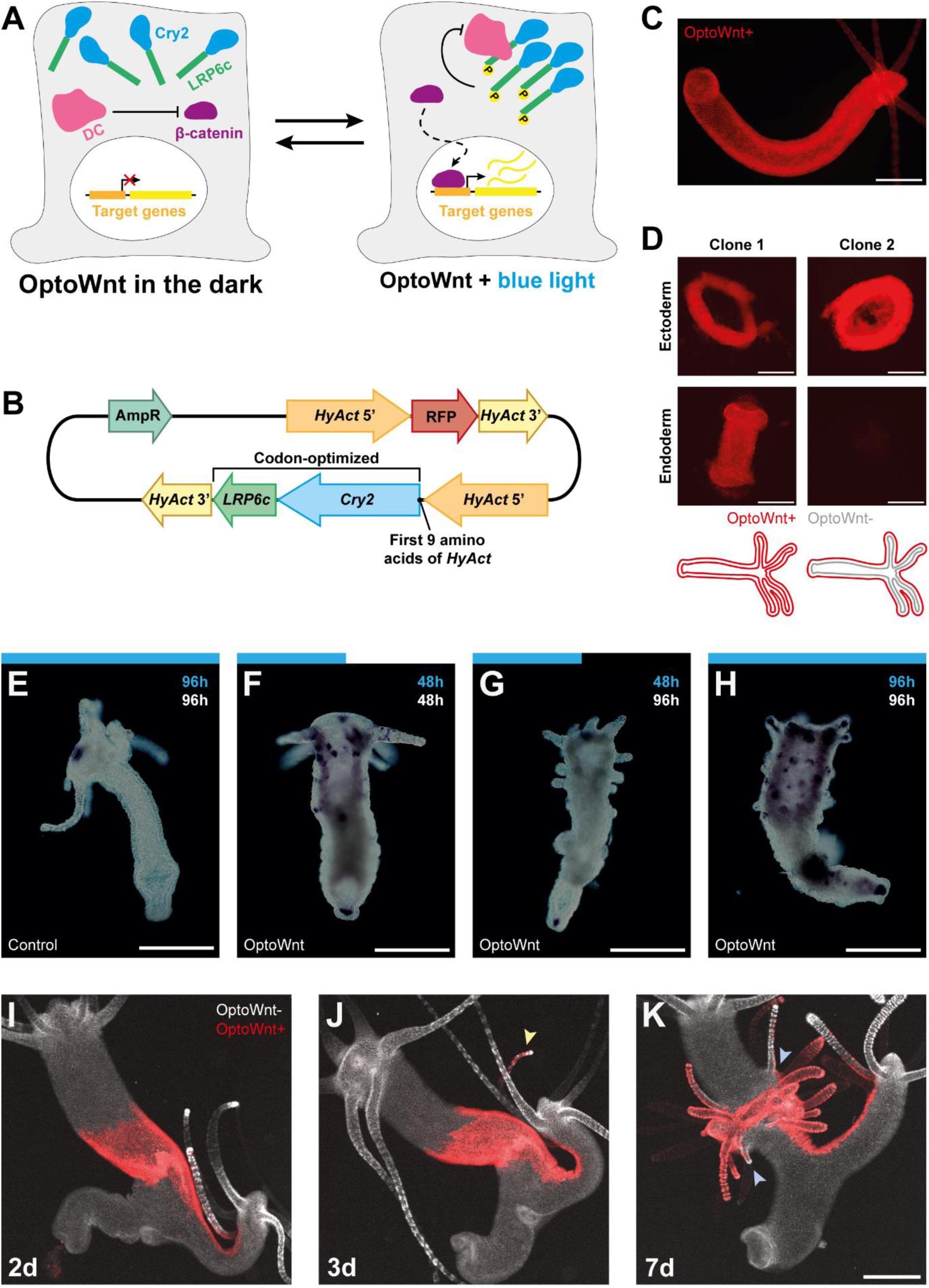
Establishment of an optogenetic system for the activation of canonical Wnt signaling in *Hydra*. (**A**) Schematic of the optoWnt system. Blue light stimulation induces Cry2-mediated clustering of the LPR6 cytoplasmic domain (LRP6c), resulting in the inhibition of the β-catenin destruction complex (DC), stabilization of β-catenin and activate transcription of Wnt target gene transcription. (**B**) Schematic of the *Hydra* optoWnt expression construct. (**C**) Representative fluorescence image showing ubiquitous RFP expression in a transgenic optoWnt *Hydra*. (**D**) Fluorescence images of separated epithelial layers from two independent optoWnt lines. Clone 1 expresses optoWnt in both ectoderm and endoderm, whereas in clone 2 optoWnt is restricted to the ectoderm. Schematic below summarizes the corresponding expression pattern. (**E–H**) Representative whole-mount in situ hybridization for *Wnt3* in control (E) and optoWnt animals (F–H) following blue-light stimulation (16 μW/mm²) (*N* = 2, for Ecto:GFP: *n* = 8, for optoWnt: *n* = 10). Control animals maintained a single endogenous *Wnt3* expression domain at the hypostome, whereas optoWnt animals developed ectopic *Wnt3* expression domains and corresponding ectopic patterning phenotypes. Light exposure is indicated by blue bars above each panel; blue numbers indicate cumulative illumination time, and white numbers indicate the time of sample collection. (**I-K**) Organizer assay using tissue grafts, where rings of optoWnt tissue (red) where grafted into GFP-expressing host animals (white) and stimulated with blue light. The samples were imaged after 2 (I), 3 (J) and 7 days (K). Ectopic tentacles first emerged after 3 days (yellow arrowhead), and by 7 days secondary head structures had formed within the graft region, incorporating both donor- and host-derived tissue (blue arrowheads), demonstrating that optogenetic activation of Wnt signaling is sufficient to induce organizer activity. Scale bars, 500 μm.

To determine whether optogenetic stimulation activates Wnt signaling *in vivo*, animals were exposed to continuous blue light (16 μW/mm^2^). Control animals lacking the optoWnt construct developed normally under identical illumination conditions and maintained a single endogenous *Wnt3* expression domain at the hypostome (Fig. 1E). In contrast, optoWnt animals exposed to blue light developed ectopic *Wnt3* expression domains along the body column and in the aboral region (Fig. 1F-H). Interestingly, the phenotypic outcome depended on the timing and duration of light exposure. Animals exposed to light for 48 h and subsequently returned to darkness developed ectopic tentacles while the ectopic *Wnt3* expression domains resolved over time (Fig. 1G). By contrast, animals maintained continuously under light for the same total duration also formed ectopic tentacles yet retained the ectopic *Wnt3* expression domains (Fig. 1H). These observations indicate that sustained optogenetic activation leads to phenotypic alterations consistent with pharmacological or genetic overactivation of Wnt signaling (12, 21, 22).

To test whether optoWnt-expressing tissue could function as an organizer, we generated chimeric animals by grafting optoWnt tissue rings into non-transgenic hosts. Upon blue light stimulation, ectopic tentacles and secondary axes emerged from the graft region (Fig. 1I-K). These secondary structures incorporated both donor-derived and host-derived tissue, demonstrating that optogenetically activated Wnt signaling is sufficient to induce ectopic patterning through the recruit of neighboring wild-type cells. Together, these findings establish optoWnt Hydra as a robust and inducible system for manipulating Wnt signaling and organizer activity during pattern formation.

### Optogenetic activation of Wnt signaling induces dose-dependent patterning responses

Under dark conditions, optoWnt animals developed largely normally, with only occasional axial abnormalities comparable to those observed in other transgenic *Hydra* lines. Continuous blue light exposure, however, induced a spectrum of phenotypes associated with ectopic Wnt signaling activation (Fig. 2A). Mild phenotypes appeared as ectopic tentacles adjacent to the endogenous head region. With increased light exposure, ectopic tentacles expanded along the upper body column, endogenous tentacles shortened, and body morphology became progressively distorted. Under the strongest effect observed, the entire body column became covered with tentacle structures. In many animals, prolonged activation additionally resulted in branched morphologies consistent with ectopic organizer formation and possibly impaired bud detachment.

**Figure 2.**
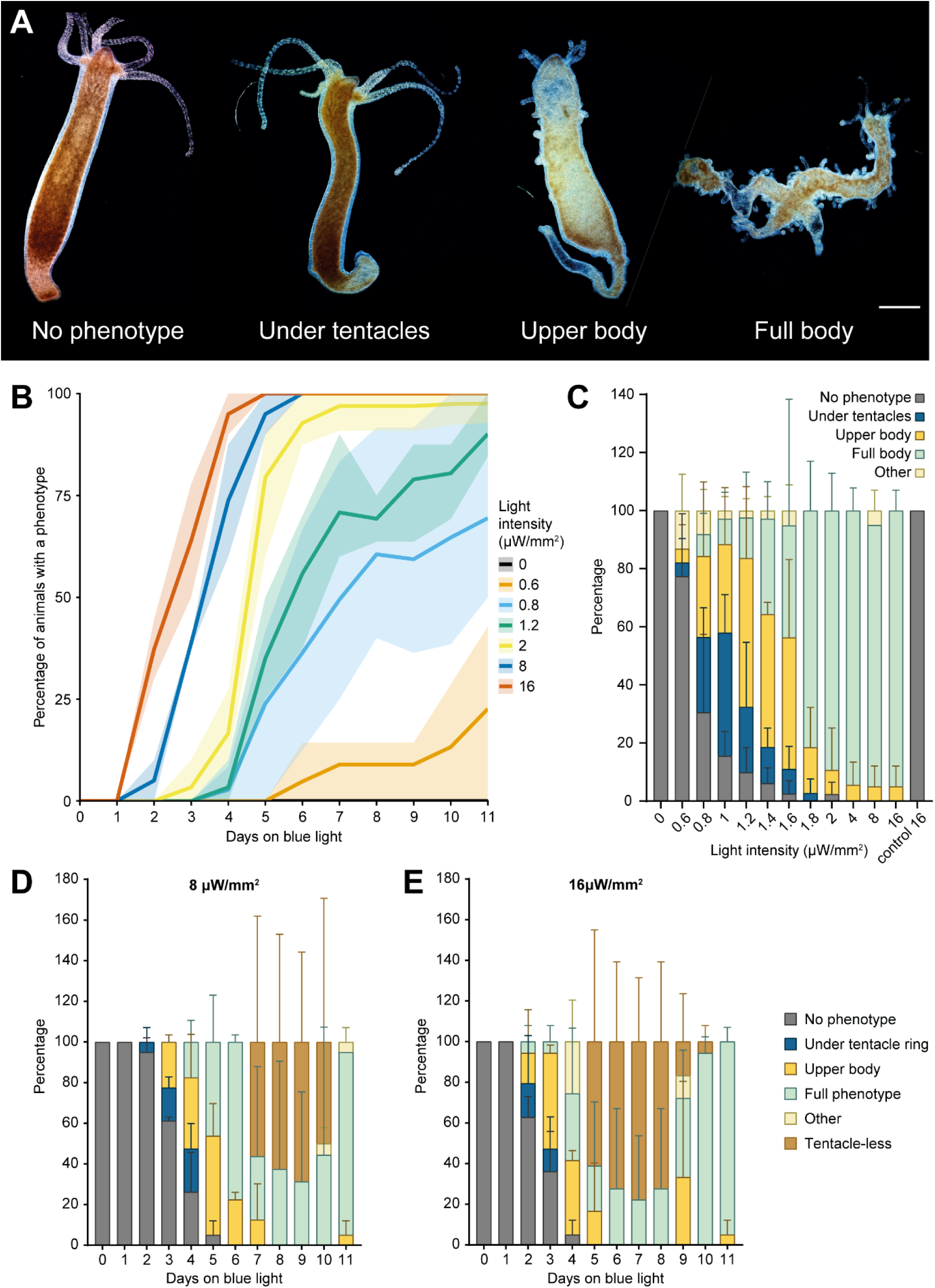
Optogenetic Wnt activation drives progressive, dose-dependent patterning responses. (**A**) Representative phenotypic stages observed during optogenetic activation of Wnt signaling, ranging from normal morphology to widespread ectopic tentacle formation covering the body column. (**B**) Fraction of animals displaying any optoWnt phenotype over time at selected light intensities. Higher light intensities accelerated phenotypic progression, whereas weaker stimulation delayed onset. Solid lines indicate the mean and shaded regions represent the min–max range. (**C**) Distribution of phenotypic stages after 11 days of continuous illumination across the tested range of light intensities. Increasing light intensity shifted the population toward progressively more advanced phenotypic states. (**D-E**) Temporal progression of phenotypic states under continuous illumination at 8 μW/mm² (D) and 16 μW/mm² (E). Animals progressed through similar phenotypic stages under both conditions, with higher light intensity primarily accelerating progression through the common sequence. Data in (C–E) are presented as mean ± SD. N = 3 independent experiments for 0–2 μW/mm² and N = 2 for 4–16 μW/mm², with n = 8–10 animals per experiment. Scale bars, 500 μm.

The emergence and progression of these phenotypes exhibited a strong dependence on light intensity. Exposure to 16 μW/mm² of blue light induced morphological changes in all animals within four days, whereas at low intensities (0.6 μW/mm²) phenotypic alterations emerged only after several days of stimulation (Fig. 2B; Suppl. Fig. 2A-E). Intermediate light intensities produced intermediate rates of phenotypic progression, indicating that Wnt pathway activation scales quantitatively with optogenetic input strength. Importantly, the distribution of phenotypic states over time suggested that stimulation intensity and exposure duration were partially interchangeable parameters. Across all tested conditions, animals progressed through similar phenotypic states in a stereotyped temporal sequence, with higher light intensities primarily accelerating progression along this trajectory (Fig. 2C-E). Stimulation with less intense light resulted in delayed onset of the phenotypic alterations but eventually arrived at the same phenotypic spectrum. This relationship suggested that cumulative signaling exposure, rather than instantaneous signaling strength, determines morphogenetic outcome.

At the highest stimulation intensities, animals transiently entered a reversible tentacle-less state characterized by reduced tissue integrity and a notable absence of tentacles (Fig. 2D,E; Suppl. Fig. 2F). Striking because ectopic tentacle formation is typically a key phenotypic hallmark of overdriven Wnt signaling. Notably, animals recovered from this state and progressed toward stable ectopic patterning phenotypes (Suppl. Fig. 2G). These observations suggest that excessively strong Wnt activation transiently disrupts tissue patterning progression, while sustained pathway activation remains compatible with organizer formation and long-term survival.

### Distinct optogenetic stimulation regimes converge onto common transcriptional trajectories

To characterize the transcriptional dynamics underlying optogenetically induced patterning responses, we performed RNA sequencing on optoWnt animals exposed to different light intensities over a temporal series spanning 48 h after stimulation onset (Fig. 3A). Principal component analysis revealed that samples separated primarily along the first principal component (PC1) according to stimulation duration and intensity, whereas variation along PC2 was largely attributable to genetic background differences (Fig. 3B,C). Notably, samples from different stimulation regimes arranged along a common trajectory in transcriptional space. In particular, samples exposed to lower light intensities for longer durations occupied similar regions of transcriptional space as samples exposed to higher light intensities for shorter periods, suggesting that distinct combinations of signal strength and duration can drive convergent transcriptional responses.

**Figure 3.**
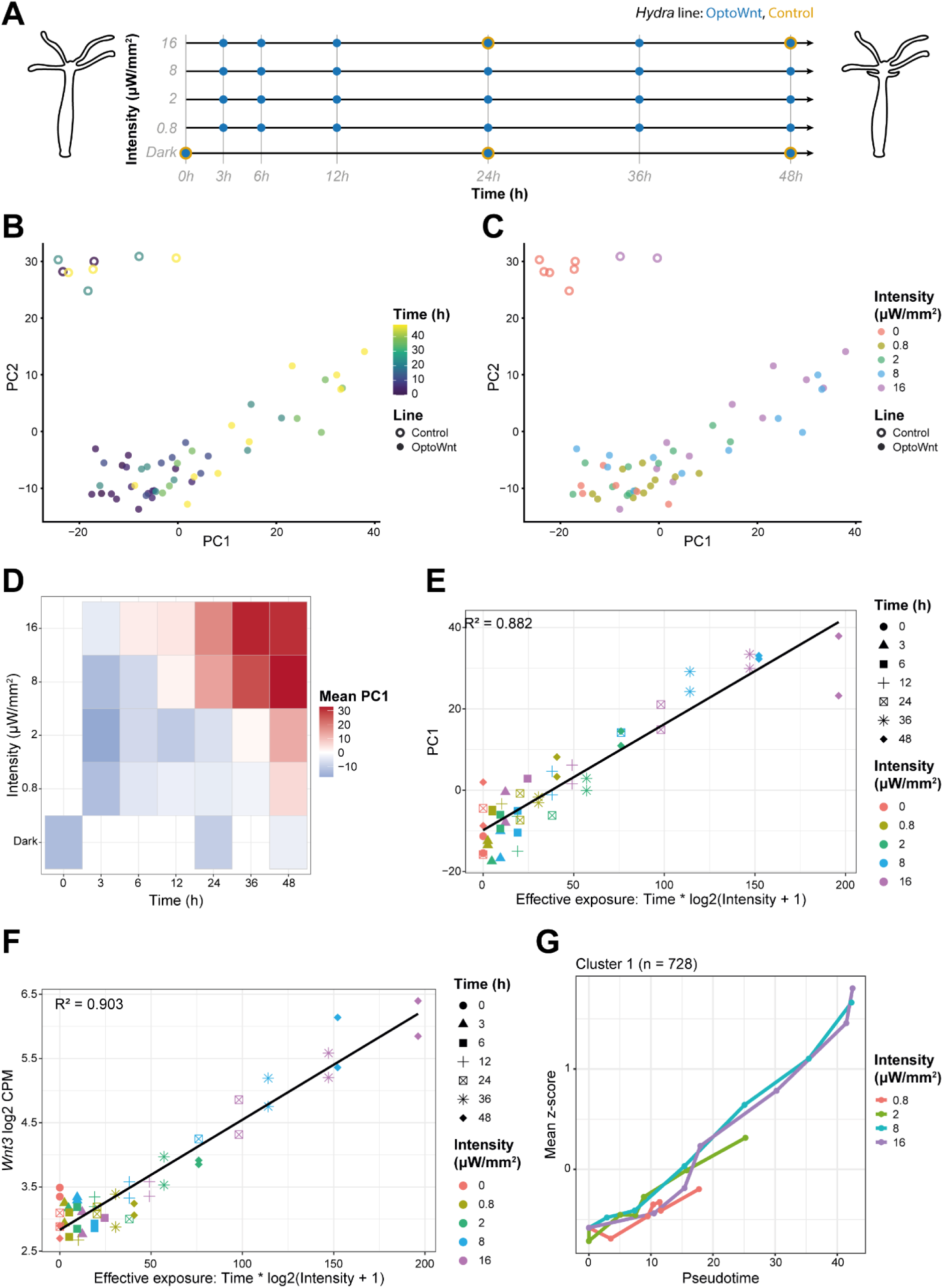
Transcriptional progression depends on the cumulative duration and intensity of optogenetic Wnt activation. (**A**) Experimental design for RNA sequencing, where optoWnt *Hydra* were exposed to different blue-light intensities over a 48h time course before RNA collection. Collection conditions are indicated by a blue dot (optoWnt *Hydra*) and orange ring (control *Hydra*). (**B,C**) Principal component analysis (PCA) of the transcriptomes colored by stimulation duration (**B**) or light intensity (**C**). (**D**) Mean PC1 values plotted as a function of stimulation duration and light intensity. (**E**) PC1 values for optoWnt samples plotted against the phenomenological effective exposure metric, defined as E_ff_ = Time * log2(Intensity + 1). Samples from different stimulation conditions fall along a linear trend (R^2^=0.882), indicating that the major transcriptional progression axis is largely determined by cumulative signaling exposure. (**F**) *Wnt3* expression (log2 CPM) plotted against effective exposure. *Wnt3* expression scales linearly with cumulative signaling exposure across stimulation conditions (R^2^=0.903). (**G**) Mean gene-wise scaled expression (z-score) of genes in Cluster 1 plotted against PCA principal curve-derived pseudotime. Samples exposed to lower intensities align with earlier portions of the trajectory generated under maximal stimulation, indicating slower progression along a common transcriptional program.

To visualize how transcriptional state varied across combinations of stimulation intensity and duration, we mapped the corresponding PC1 values onto a two-dimensional intensity-duration grid (Fig. 3D). Strikingly, samples with similar PC1 values aligned diagonally, linking early time points under strong stimulation to later time points under weaker stimulation. A similar diagonal organization was observed for numerous Wnt-responsive genes, including *Wnt3*, *Bra1*, *Axin*, and *Sp5* (Suppl. Fig. 3). These observations suggest that comparable transcriptional outputs can be achieved through different combinations of signal intensity and exposure duration.

To identify a quantitative descriptor that could account for the transcriptional responses induced by different combinations of stimulation intensity (I) and duration (t), we sought an effective exposure (E_eff_) metric directly from the data. At fixed light intensity, the expression of many Wnt-responsive genes increased approximately linearly with time (Suppl. Fig. 4). However, the rate of this increase depended nonlinearly on light intensity and exhibited apparent saturation. Empirically, these rates were approximately linear with the logarithm of light intensity (Suppl. Fig. 4). These observations motivated the definition of a phenomenological effective exposure metric proportional to the product of stimulation duration and the logarithm of light intensity (E_eff_ ∝ t*log(I+1)). When PC1 values were plotted against this metric, samples from all stimulation conditions collapsed onto a common linear trajectory (Fig. 3E), indicating that transcriptional state is largely determined by cumulative signaling input. *Wnt3* expression levels showed a similarly strong relationship with effective exposure across all stimulation conditions (Fig. 3F). Thus, effective exposure provides a low-dimensional description of how signal magnitude and duration combine to shape Wnt-induced transcriptional responses.

Differential expression analysis identified six gene clusters with temporally ordered responses downstream of optogenetic Wnt activation (Suppl. Fig. 5). Across several of these clusters, transcriptional responses progressed more slowly under lower-intensity stimulation than under stronger stimulation. To quantify this relationship, we ordered samples along a PCA-derived pseudotime trajectory representing progression through the common transcriptional state space (Suppl. Fig. 6). When the mean expression of genes in the largest cluster was plotted against pseudotime, samples exposed to lower light intensities aligned with earlier portions of the trajectory generated under maximal stimulation (16 μW/mm²) (Fig. 3G). A similar relationship was observed across the remaining major clusters (Suppl. Fig. 7), together comprising more than three-quarters of all differentially expressed genes. These findings indicate that distinct combinations of stimulation intensity and duration drive tissues along the same transcriptional trajectory, with cumulative signaling exposure determining the extent of transcriptional progression. They further predict that equivalent morphogenetic outcomes should be achieved once tissues reach comparable transcriptional states, irrespective of how that cumulative signaling exposure was attained.

### Successful head regeneration depends on cumulative Wnt signaling exposure

To directly test whether cumulative Wnt signaling determines morphogenetic outcome, we used epithelial spheroid regeneration assays under isotonic conditions, which normally inhibit head regeneration by preventing the mechanical oscillations required for organizer formation (21, 23). Under normal freshwater conditions, small fragments of *Hydra* tissue fold into spheroids and regenerate complete animals with a correctly patterned oral–aboral axis (Fig. 4A). Consistent with previous studies, spheroids regenerating under isotonic conditions failed to form heads and remained spherical (Fig. 4B). By contrast, optoWnt spheroids exposed to blue light successfully regenerated and developed head structures despite the isotonic conditions (Fig. 4C).

**Figure 4.**
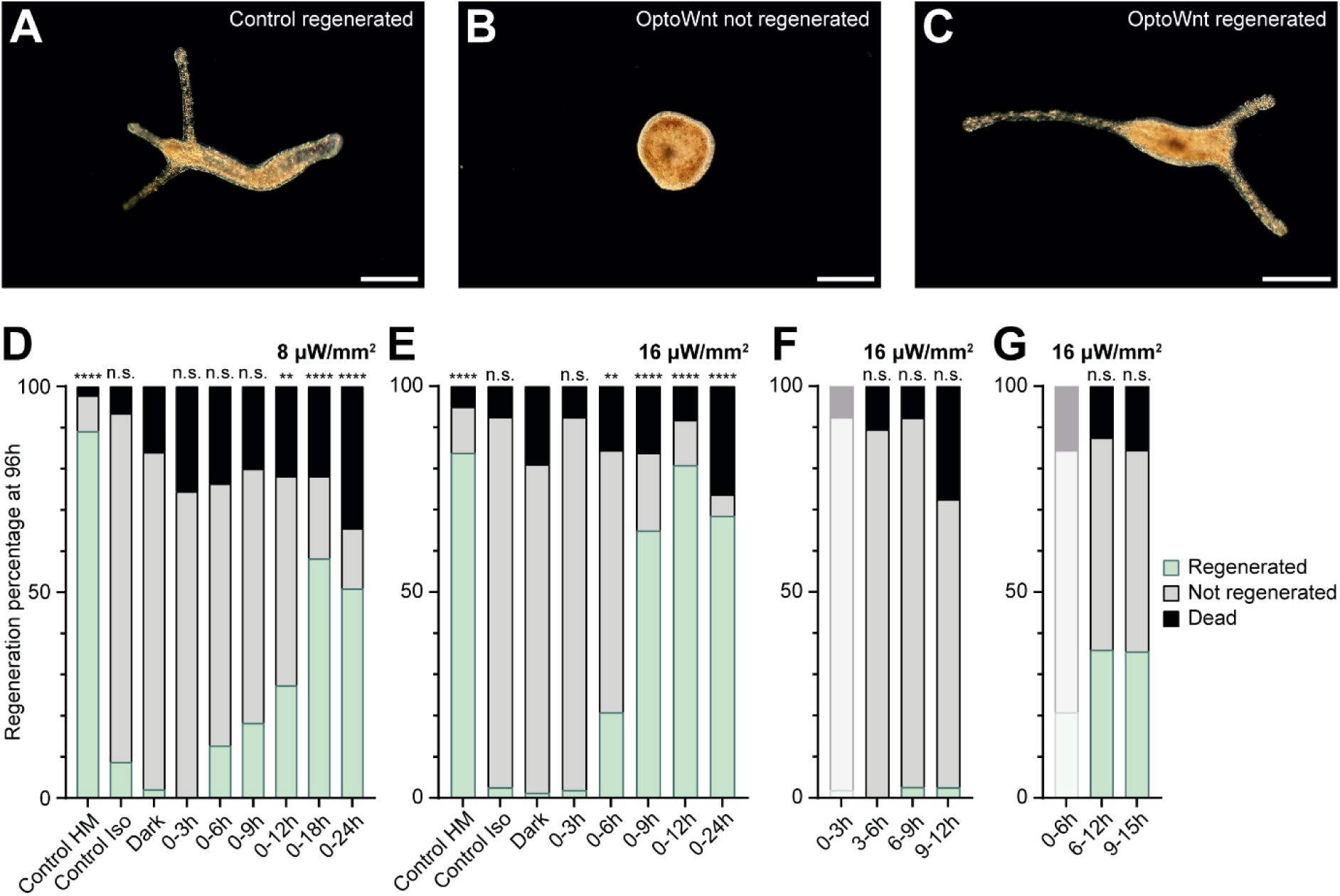
Cumulative Wnt signaling, rather than a discrete temporal window, determines successful regeneration. (**A–C**) Representative spheroid regeneration outcomes after 96 h. Under normal freshwater conditions, epithelial spheroids regenerate complete animals with a correctly patterned oral–aboral axis (A). Under isotonic conditions (70 mM sucrose), regeneration was inhibited and spheroids failed to form head structures (B). Optogenetic activation of Wnt signaling rescued regeneration under isotonic conditions, resulting in successful head formation (C). (**D-E**) Regeneration outcomes following different durations of optogenetic stimulation at 8 μW/mm² (D) or 16 μW/mm² (E). Increasing stimulation intensity reduced the duration of illumination required to rescue regeneration, indicating that signal strength and exposure time act as partially compensatory parameters. Regeneration was scored after 96 h. The first two columns show reference distributions for successful regeneration in *Hydra* medium (Control HM) and unsuccessful regeneration under isotonic conditions (Control Iso), both using the Ecto::GFP control line. Statistical comparisons are shown relative to optoWnt spheroids regenerating under isotonic conditions in the absence of blue-light stimulation. (**F-G**) Regeneration outcomes following optogenetic stimulation restricted to 3 h (F) or 6 h (G) sliding windows during regeneration. Regeneration efficiency depended primarily on the total duration of Wnt activation rather than the developmental timing of stimulation, arguing against the existence of a narrow temporal competence window for organizer formation. Data in (D–G) were analyzed using χ² tests with Holm–Bonferroni correction. For (D), N = 2 independent experiments with total n ≥ 46 per condition; for (E–G), N = 2–4 independent experiments with total n ≥ 38 per condition. \*\**P_adj_* < 0.01 \*\*\*\**P_adj_* < 0.0001. n.s., not significant. Scale bars, 250 μm.

Regeneration success depended strongly on both stimulation intensity and exposure duration. At 8 μW/mm², successful regeneration required at least 12–18 h of stimulation, whereas shorter exposures failed to rescue head formation (Fig. 4D). Increasing the light intensity to 16 μW/mm² reduced the required stimulation duration to 6–9 h to produce comparable regeneration efficiencies (Fig. 4E). Thus, stronger stimulation could partially substitute for longer exposure, indicating that signal magnitude and duration behave as compensatory parameters during organizer formation. These findings are consistent with the transcriptional analyses and suggest that morphogenetic outcomes are governed by cumulative Wnt signaling exposure rather than instantaneous pathway activity.

To determine whether regeneration depends on signaling during a specific temporal window, we next restricted optogenetic stimulation to sliding windows of either 3 or 6 h at different stages of regeneration (Fig. 4F,G). Regardless of when stimulation was applied, 3 h windows were insufficient to rescue regeneration. Six-hour stimulation windows modestly increased regeneration efficiency but remained unable to restore regeneration to levels achieved by longer durations. Ultimately, regeneration outcomes depended primarily on the total duration of Wnt activation rather than on the developmental timing of stimulation. Together, these findings demonstrate that organizer formation is determined by the integrated history of Wnt signaling rather than activation during a discrete competence window.

## Discussion

Wnt signaling is a central regulator of developmental patterning and regeneration (24, 25), yet how cells interpret dynamic pathway activity to generate stable morphogenetic outcomes remains an open question. Here, using the first optogenetically activatable *Hydra* line, we show that successful organizer formation depends on cumulative Wnt signaling exposure rather than activation within a narrowly defined temporal window. By systematically varying both the duration and intensity of pathway activation, we demonstrate that morphogenetic outcomes arise through temporal integration of signaling input, revealing that signal strength and exposure time behave as partially compensatory parameters. Beyond establishing optogenetics as a powerful new experimental approach in *Hydra*, our findings suggest that tissues decode morphogen signals by integrating cumulative pathway activity over time and progressively accumulating transcriptional responses until organizer-forming thresholds are reached.

OptoWnt activation in *Hydra* required substantially higher light intensities than previously reported in mammalian systems (20), potentially reflecting differences in the tissue’s optical properties, expression levels, or Cry2 dynamics. Nevertheless, prolonged illumination alone produced no detectable defects in control animals, demonstrating that the system is well-suited for long-term *in vivo* perturbation experiments. Importantly, adapting the optoWnt construct required only minimal modifications to the original mammalian system, suggesting that many existing optogenetic tools may be readily transferable to *Hydra*. Beyond enabling precise temporal control of signaling activity, optogenetics opens new opportunities to interrogate spatial aspects of pattern formation, including organizer induction, axis competition, and scaling mechanisms during regeneration. Together, these results establish *Hydra* as a tractable system for quantitatively dissecting how dynamic signaling inputs regulate developmental patterning *in vivo*.

Phenotypically, optoWnt activation produced animals resembling a combination of the phenotypes associated with *Wnt3* overexpression and pharmacological activation of canonical Wnt signaling by alsterpaullone (AP) (12, 26). Similar to AP treatment, optoWnt acts downstream of ligand-receptor interactions by stabilizing β-catenin signaling output. However, unlike pharmacological perturbations, optogenetics allows the magnitude and duration of pathway activation to be manipulated independently and reversibly (17), enabling direct interrogation of how tissues decode dynamic Wnt inputs. Using this approach, we show that distinct combinations of signal intensity and duration can produce equivalent transcriptional and morphogenetic outcomes. *Hydra* exposed transiently to AP often require several additional days to develop ectopic tentacles after drug removal (26). Our results therefore raise the possibility that part of the delay observed after AP exposure reflects drug persistence rather than only the intrinsic timescale of morphogenetic commitment itself. Once a transcriptional threshold compatible with organizer formation has been reached, morphological progression appears to proceed relatively rapidly.

One particularly striking observation was the transient tentacle-less state induced by sustained high-intensity stimulation. During this phase, animals displayed contracted morphologies and reduced tissue integrity before subsequently recovering and developing stable ectopic tentacle phenotypes. We speculate that excessive optogenetic activation initially drives Wnt signaling beyond its normal physiological range, resulting in a temporary disruption of tissue homeostasis. The eventual recovery of these animals nevertheless suggests that *Hydra* tissues actively buffer supraphysiological Wnt signaling and can recover and re-enter normal patterning programs following transient perturbation. This interpretation is consistent with previous observations that canonical Wnt signaling induces the expression of multiple pathway inhibitors, including negative feedback regulators such as Axin and Dkk family members, as well as Sp5, a key negative regulator of Wnt signaling specifically in the context of *Hydra* regeneration (27-29). Such feedback may contribute both to signaling robustness and to the stabilization of long-term patterning outcomes.

A central conclusion emerging from our experiments is that *Hydra* tissues interpret Wnt signaling cumulatively over time. Across transcriptional and morphogenetic readouts, stimulation intensity and duration behaved as partially interchangeable parameters. Distinct stimulation regimes converged onto common transcriptional trajectories, and successful regeneration could be induced either through strong short-term activation or weaker prolonged activation. Mechanistically, several processes may contribute to the temporal integration downstream of Wnt signaling. Progressive accumulation of β-catenin, positive feedback through endogenous Wnt ligands such as Wnt3, chromatin remodeling, or transcriptional memory mechanisms may all enable signaling history to be retained over time.

Optogenetic Wnt perturbations in mammalian stem cells have previously demonstrated how signaling heterogeneity influences developmental patterning (20, 30), while related optogenetic systems in *Xenopus* embryos have shown that ectopic Wnt activation can induce secondary organizers and axis duplication (31). More broadly, β-catenin signaling dynamics have been shown to regulate fate choice in neuronal differentiation (32, 33), and Wnt responses can shift from transient to sustained depending on developmental context and pathway crosstalk (34). By contrast, in fibroblast reprogramming, the timing of pathway activation rather than its duration appears to be the dominant determinant of cellular outcome (35). These observations suggest that signaling dynamics are interpreted through context-dependent decoding strategies that vary between developmental systems.

Our findings also contribute to a broader understanding of how morphogen signaling dynamics are decoded across biological systems. In the classical view, cells acquire positional information by interpreting morphogen concentration thresholds (36-38). Increasing evidence, however, indicates that developmental decisions depend not only on morphogen concentration but also on the duration and cumulative history of signaling activity (37-39). In other systems, Bmp (40, 41), Sonic Hedgehog (42), Nodal (43), ERK (44), and Wnt signaling (30, 32) have all been shown to exhibit duration-dependent or cumulative effects on fate specification. Our results extend these concepts to *Hydra* regeneration and suggest that the temporal integral of signaling activity can itself constitute an instructive variable. More broadly, temporal integration may represent a conserved strategy through which tissues buffer fluctuating signaling inputs while maintaining robust patterning outcomes.

Together, our findings support a model in which organizer formation is governed by the integrated history of Wnt signaling rather than instantaneous pathway activity. More broadly, our work suggests that developmental systems can decode dynamic morphogen inputs through temporal integration of cumulative signaling exposure, progressively converting transient signaling events into stable patterning decisions.

## Materials & methods

### *Hydra* culture conditions

All procedures were carried out in compliance with ethical guidelines of Swiss national regulations for animal research. *Hydra* were kept in Volvic water at 18°C and fed freshly hatched *Artemia nauplii* three times per week. OptoWnt *Hydra* were cultured in the dark to prevent premature phenotype formation from the ambient light. For experiments, only animals without sexual organs, buds and a phenotype were selected. Unless otherwise specified, animals expressing green fluorescent protein (GFP) in the ectoderm (Ecto::GFP, control) (45) and optoWnt clone 1 (positive in both epithelial layers) were used.

### OptoWnt plasmid assembly and clonal line generation

The codon-optimized Cry2-LRP6c sequence, including the first 27 base pairs of *HyAct*, was commercially produced (IDT, gblock). The plasmid backbone AR-ActpA, a derivative from the pBSAA-AR cloning vector with *HyAct* 5’ and 3’ sequences flanking an unoccupied cloning site, was digested with BamHI and XbaI (Suppl. Fig. 1A)(46). After digest purification, the digested plasmid and optoWnt DNA fragment were assembled using Gibson assembly. The resulting construct (9901 bp) was propagated in ultracompetent cells from the XL10-Gold strain and purification was performed using the commercially available Qiagen QIAfilter Plasmid Midi kit (Qiagen, 12243). The purified DNA was sequenced to confirm sequence integration had occurred as expected (primer sequences in Suppl. Table 1) (Suppl. Fig 1B).

The optoWnt construct was microinjected into fertilized embryos of *Hydra vulgaris* strain AEP (clone F2A4 Kiel) as previously described (47). Hatchlings were fed manually until they had grown enough to eat independently. RFP+ hatchlings were left to develop, while non-fluorescent animals were removed. The *Hydra* cultures were constantly monitored and only animals with the highest percentage of RFP+ cells were kept. Once a handful of animals from each clone had a significant enough patch of red cells, they were exposed to 0.8 μW/mm^2^ blue light to examine whether they responded to the blue light. Only clones of which an animal developed a phenotype were selected. For each of the remaining clones, a single, completely RFP+, animal was once more used to re-establish the clonal lines.

### Separating *Hydra*’s epithelial layers

The protocol for the separation of the ectodermal and endodermal layers was adapted from a protocol for the generation of recombined *Hydra* (48). Solutions A and B were prepared containing equal volumes of 1% Procaine-HCl in MilliQ, *Hydra* medium (HM; 1 mM CaCl2, 0.2 mM NaHCO3, 0.02 mM KCl, 0.02 mM MgCl2, and 0.2 mM tris-HCl (pH 7.4)) and dissociation medium (DM; 3.6 mM KCl, 6 mM CaCl2, 1.2 mM MgSO4, 6 mM sodium citrate, 6 mM sodium pyruvate, 4 mM glucose, and 12.5 mM N-tris(hydroxymethyl)methyl-2-aminoethanesulfonic acid (pH 6.9)) with pH 4.5 and 2.5, respectively. For ectoderm retrieval, *Hydra* were incubated in solution A for 5 minutes at 4°C followed by 1.5 minutes at in solution B at 4°C. Endoderm was prepared by 5 minutes in solution A at 4°C followed by 3 minutes in solution B at 4°C. After the incubations, all tissues were kept in DM at 18°C for imaging.

### Optogenetic activation

For optogenetic stimulation, *Hydra* were moved to black-walled 24-wells plates (IBIDI, 82426) and placed onto a LAVA illumination device (20, 49, 50). In short, the illumination device is driven by a Raspberry Pi computer and allows for the programming of custom illumination patterns. Each of the 24 wells can be illuminated independently. Blue light was produced by 470 nm LEDs.

### Localized optoWnt activation in chimeric animals

Glass needles were pulled from glass capillaries with a 1.2 mm outer diameter (Harvard Apparatus, 30-0050). To create chimeras, a ring of tissue was cut from the middle of the body column of optoWnt *Hydra*. A similar tissue ring was removed from *Hydra* expressing GFP-tagged β-catenin and replaced by the optoWnt tissue (46). The three tissue pieces were slid onto a glass needle followed by a small piece of parafilm to hold the tissue in place. After 2-3h the now healed animals were removed from the glass needle and left to recover in HM for five days. On the last day of this recovery period, the animals were fed.

The animals were exposed to 0.8 μM/mm^2^ blue light for 7 days. During this time phenotype development was monitored and representative images were taken. GFP and RFP were used to identify the background and optoWnt tissue, respectively.

### Scoring optoWnt phenotypes

*Hydra* exposed to continuous illumination were fed only once per week but monitored daily. For clarity, phenotypes were divided into four distinct categories: ‘no phenotype’, ‘under tentacles’, ‘upper body’ and ‘full body’. If an animal did not fall into any of these categories, it was labelled as ‘other’. Animals exposed to high intensity light went through a tentacle-less phase where the phenotypic stage could not be assigned reliably, these animals were labelled as ‘overexposed’.

### Spheroid preparation

Spheroids were prepared following an established protocol (51). In brief, animals were bisected in the middle of the body column. From each half, a tissue ring was obtained through a subsequent, parallel cut. These rings were divided into two to four rectangular pieces, depending on their size. Tissue fragments were left to fold into spheroids for three to four hours in DM at room temperature. Spheroids of typical size (250-350 μm) and that had closed properly were selected for the experiments.

### Scoring spheroid regeneration

For all conditions, HM was supplemented with kanamycin (50 μg/ml) and streptomycin (100 μg/ml). For sucrose conditions, sucrose was dissolved to a concentration of 70mM. Before use, the medium was filter sterilized. Successfully folded spheroids were transferred to antibiotic-supplemented HM and randomly distributed across the wells of a black-walled 24-well plate (IBIDI, 82426) in a minimal volume of medium. Medium with or without sucrose was added to each well, then the plate was moved onto the illumination device. For every condition, an intensity of 16 μW/mm^2^ was applied, time windows were programmed in the associated software. Regeneration was scored as ‘regenerated’, ‘not regenerated’ or ‘dead’ after 96 hours.

For statistical comparison, the R package ‘rstatix’ was used (52). A contingency table showing the number of spheroids per category for each sample was created. A chi-squared test on the full contingency table indicated significant differences between the samples. Pairwise comparisons were subsequently performed to identify differences between individual samples, a p-value correction was performed using the Holm-Bonferroni method (36 total comparisons for 8 μW/mm^2^ & 78 total comparisons for 16 μW/mm^2^).

### Imaging

Unless otherwise specified, images were acquired using the Revolve hybrid microscope from Echo (RVL2-K), using the inverted mode with a 4.0×/0.16 UPlanXApo objective (Olympus, AMEP4904). A transmission light camera (CMOS-color) was used for dark-field images, while a black/white camera (sCMOS-monochrome) was used for fluorescence.

Grafted *Hydra* were imaged using a Zeiss Axio Observer 7 microscope equipped with a CSU-W1 spinning disk confocal scanning unit (Yokogawa) featuring a 50-μm pinhole disk, a 5×/0.25 FLUAR objective (Zeiss, 440125-0000-000), and two Prime 95B scientific complementary metal-oxide semiconductor (sCMOS) cameras (Photometrics). VisiView imaging software (Visitron) was used to control the system. At each timepoint, a single plane was acquired using both the 488 and 561 nm lasers at a 1200 by 1200 pixel resolution with a 2.2-μm pixel size.

### RNA sequencing of the early Wnt response in optoWnt *Hydra*

To investigate the early Wnt response, a variety of timepoints spread across 48 hours and intensities ranging from no light to 16 μW/mm2 were included. Per duplicated sample, 10 *Hydra* were collected, immediately homogenized in TRI reagent® (Sigma, T9424) and stored at -80°C until further use. RNA extraction was performed using the commercially available Zymo Direct-zol RNA mircroPrep kit (Zymo, R2060). Centrifugation times were doubled to ensure complete flow-through of the columns.

RNA integrity was assessed using the Agilent TapeStation system. Then, mRNA-seq libraries were prepared using the Illumina Stranded mRNA Prep kit according to the manufacturer’s instructions. The libraries were sequenced on an Illumina NovaSeq 6000 platform using paired-end 56 bp reads, with demultiplexing performed using bcl2fastq2.

### Analysis of Optogenetics Bulk RNA-seq

Paired-end RNA-seq reads were aligned to the *Hydra vulgaris* AEP genome assembly HydraT2T_AEP (NCBI RefSeq GCF_038396675.1), and gene-level counts were generated with STAR v2.7.10b using --quantMode GeneCounts and the matching processed GTF annotation. For preliminary sample quality control gene counts from all sequenced samples were library normalized to obtain CPM and log2 transformed CPM values using a pseudocount of 8.

Sample quality was assessed by correlating each sample with its exact replicate and with a neighborhood pseudo-sample formed from nearby conditions of the same line. For controls, the neighborhood consisted of all other control samples. Samples with the lowest neighborhood correlation were flagged as low quality and used to identify outliers for exclusion.

For downstream analyses, gene counts from samples retained after quality control were normalized with edgeR using TMM normalization and converted to log2 CPM values with a prior count of 8. For exploratory analyses and visualization, informative genes were selected using non-control (optoWnt) samples only. Informative features were selected for significant levels of expression and variance: genes were first required to have a 95th-percentile expression greater than 8 CPM. For these expressed genes, the variance across samples was compared with the expected variance given mean expression, estimated using limma’s empirical Bayes mean–variance trend. Genes with more the 1.5 fold excess variance were retained for the final feature matrix.

Principal component analysis was performed on the centered, unscaled logCPM matrix of informative genes. Unless otherwise indicated, PCA was performed on the filtered analyzed samples including controls. PC1 was oriented so that it increased with effective exposure in optoWnt samples. For the construction of Time/Intensity grids, optoWnt samples were grouped by stimulation duration and intensity, and the mean PC1 or specific gene expression values were calculated for each observed condition. Linear fits of PC1 or *Wnt3* expression against effective exposure were performed using optoWnt samples only. Effective exposure was defined as Time × log2(Intensity + 1).

To model transcriptional responses as a function of stimulation duration and intensity, gene expression models were fitted using limma with a natural-spline model including smooth effects for time, log2-transformed intensity, and their interaction. Time was modelled with a natural spline with 3 degrees of freedom, and log2(Intensity + 1) was with a natural spline with 2 degrees of freedom. The spline knots and boundary knots estimated from the observed samples were used for prediction on the observed time-by-intensity grid. F-tests across the non-intercept model coefficients were used to identify genes with time/intensity-associated expression changes. For clustering, fitted expression values were predicted on the observed time-by-intensity grid, z-scored gene-wise across this grid, and genes significant at FDR < 0.01 were clustered by k-means into six clusters (nstart = 50). Clusters were renumbered by decreasing cluster size.

Pseudotime was derived from the PCA representation of optoWnt samples. A principal curve was fitted through the PC1–PC2 coordinates using princurve::principal_curve with smoother = "smooth_spline", stretch = 10000, and thresh = 0.1. The resulting curve position was used as PCA pseudotime, oriented to increase with effective exposure, and scaled to the 0–48h range for plotting. Gene and cluster trajectories were plotted against PCA pseudotime after averaging replicates by intensity and time, with the shared Dark/0 h baseline added to each non-zero intensity trajectory. For cluster-level trajectories, expression values were first scaled gene-wise across samples and subsequently averaged across all genes in the cluster.

## Acknowledgements

We would like to thank the members of the FMI Functional Genomics Facility for performing the library preparation and RNA sequencing and the Facility for Advanced Imaging and Microscopy at FMI for their support.

## Author contributions

S.L.W. and C.D.T. conceptualized the project and wrote the original draft. S.L.W., N.R., J.F., S.S., J.W., and A.K. performed experiments. S.L.W., P.P. and C.D.T. performed bioinformatic analysis. S.L.W. visualized the data. C.D.T. supervised the project.

